# Clustering and correlations: Inferring resilience from spatial patterns in ecosystems

**DOI:** 10.1101/233429

**Authors:** Sumithra Sankaran, Sabiha Majumder, Ashwin Viswanathan, Vishwesha Guttal

**Affiliations:** Centre for Ecological Sciences, Indian Institute of Science, Bengaluru, India, 560012; Department of Physics, Indian Institute of Science, Bengaluru, India, 560012; Institute of Integrative Biology, ETH Zurich, Switzerland, 8092; Nature Conservation Foundation, Bengaluru, India, 560097

## Abstract

1. In diverse ecosystems, organisms cluster together in such a manner that the frequency distribution of cluster sizes is a power-law function. Spatially-explicit models of ecosystems suggest that *loss of such power-law clustering* may indicate loss of ecosystem resilience. Hence, it is hypothesised that spatial clustering properties in ecosystems - which can be readily measured using remotely-sensed high-resolution data - can help infer proximity to ecosystem thresholds and may even provide early warning signals of ecosystem collapse. Recent empirical and simulation studies, however, don’t find consistent relationships between spatial clustering and ecosystem resilience. Furthermore, how spatial clustering metrics relate to other well-known early warning signals of ecosystems collapse, specifically the phenomenon of *critical slowing down* (CSD), remains unclear.
2. We synthesize the literature on cluster sizes in empirical and theoretical studies that show how local interactions (especially, positive feedback) among organisms can cause power-law clustering. In addition, we analyse a minimal spatial model of ecosystem transitions that allows us to disentangle the role of environmental stressor and positive feedback on spatial patterns and ecosystem resilience.
3. Our literature synthesis reveals that empirically observed power-law clustering in ecosystems is parsimoniously explained by local positive feedback. Our synthesis together with model analysis demonstrates that, depending on the strength of positive feedback, emergence of power-law clustering can occur at any distance from the critical threshold of ecosystem collapse. In fact, we find that for systems with strong positive feedbacks, which are most likely to exhibit abrupt transitions, loss of power-law clustering may not even occur prior to ecosystem thresholds. We also argue that cluster-size distributions are unrelated to the phenomenon of CSD.
4. We demonstrate that, due to CSD, a power-law feature does occur near critical thresholds but in a different quantity; specifically, a power-law decay of spatial correlations of ecosystem state.
5. We conclude that loss of power-law clustering cannot be used as a reliable indicator of ecosystem resilience. Our synthesis and model analyses highlights links between local positive feedback, emergent spatial properties and how they may be used to interpret ecosystem resilience.

## II. INTRODUCTION

Desertification of semi-arid ecosystems (van de Koppel et al. 2002), eutrophication of lakes (Carpenter et al. 1999), spread of diseases (Chaves et al. 2012), invasion (Hansen et al. 2013) and community shifts in coral reefs (Knowlton 2004) are some examples of state transitions or regime shifts in ecological systems. Some of these transitions can be abrupt and irreversible, leading to catastrophic loss of wildlife, habitats, and ecosystem services. Such transitions are also known as critical transitions in the ecology literature. They happen when a system crosses a certain threshold, called critical threshold, of environmental conditions. Over the last decade, several studies have devised and validated methods to detect the vulnerability of ecosystems to transitions (Carpenter et al. 2011, Dakos et al. 2012, 2011, Eby et al. 2017, Guttal and Jayaprakash 2009, Kéfi et al. 2014, Kéfi et al. 2007, Scheffer et al. 2009). One such method is based on the idea that patterns of self-organisation in ecosystems can offer signatures of resilience (Kéfi et al. 2014, Kéfi et al. 2007, Rietkerk et al. 2004, von Hardenberg et al. 2001). Self-organised patterns themselves often result from an interplay of facilitative and competitive interactions among organisms (Manor and Shnerb 2008, Scanlon et al. 2007, von Hardenberg et al. 2010). Therefore, a comprehensive understanding of how local interactions between organisms scale to their spatial distribution and affect ecosystem resilience, is of broad ecological interest.

Of many varieties of self-organization found in nature (D’Odorico et al. 2012, Kéfi et al. 2007, Rietkerk and van de Koppel 2008, Scanlon et al. 2007), we focus on spatial patterns where organisms exhibit clustering of irregular size and shape (see Glossary); these are found in many ecosystems such as semi-arid ecosystems, mussel beds or seagrass (Fig 1). Here, the frequency distributions of these cluster-sizes may follow a power-law function (henceforth referred to as *power-law clustering*). These are interesting because they may imply that systems lack characteristic size/shape (see Box 1 for a summary of properties of power-laws). Some simulation and empirical studies suggest that when ecosystems are stressed, clusters fragment leading to loss of large patches (Kéfi et al. 2014, Kéfi et al. 2007). This results in a qualitative change in the properties of cluster sizes, from a power-law to an exponential distribution. The progressive truncation of the tail of the power-law clustering has, therefore, been hypothesised to represent loss of resilience in ecosystems (Fernández and Fort 2009, Kéfi et al. 2014, Kéfi et al. 2007, Kéfi et al. 2011, Lin et al. 2010, Weerman et al. 2012).

**Figure 1:**
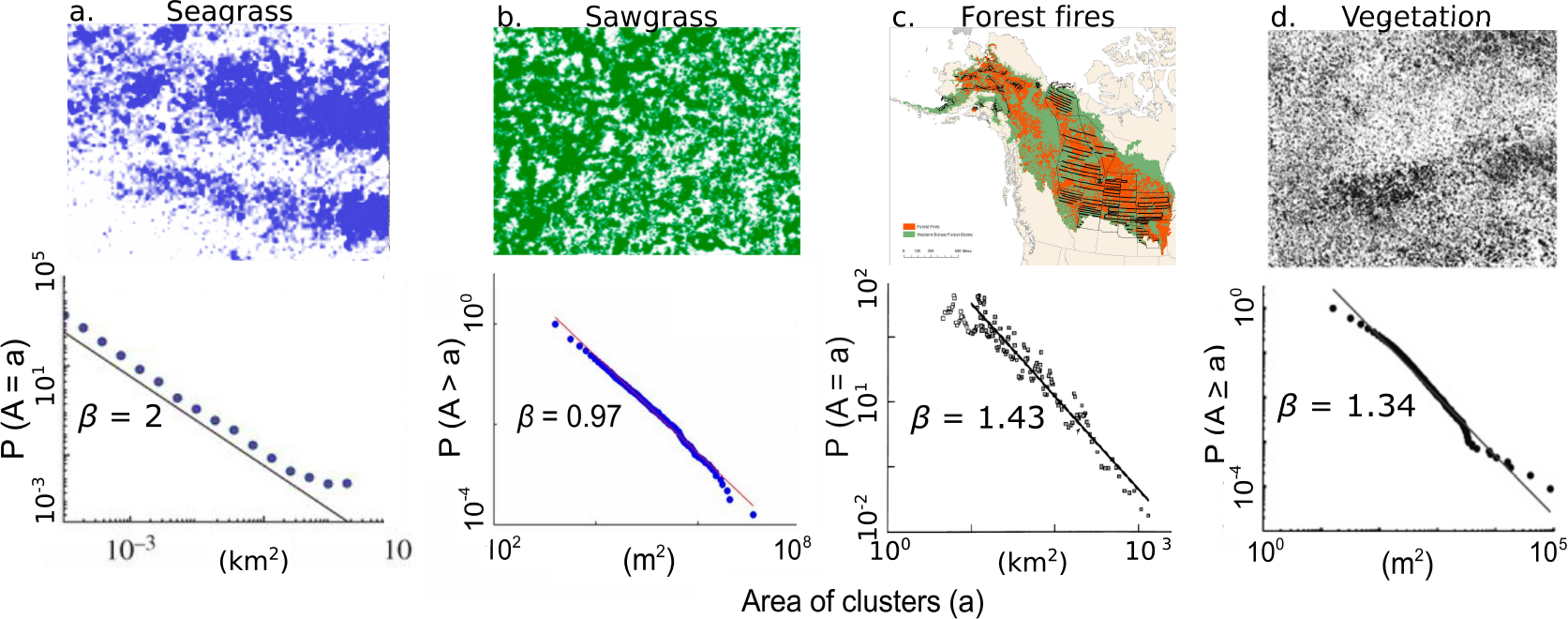
Power-law cluster size distributions in different ecosystems (top panel) and representative snapshots which are not necessarily from the same study area or time period (bottom panel). (a) West broad ledges seagrass near the isle of scilly (Irvine et al. 2016) (b) Saw-grass in everglades wetlands, USA (Foti et al. 2013) (c) Forest fires in Alaskan boreal forests, USA (1990-91) (Malamud et al. 1998) and (d) Vegetation in Kalahari, Namibia (Scanlon et al. 2007). Top row image credits: (a) modified from (Duffy et al. 2017), (b) https://doi.org/10.1016/j.ecss.2017.11.001 [CC BY] http://creativecommons.org/licenses/by/4.0/, (Foti et al. 2013), (c) U.S. Geological Survey, Department of the Interior/USGS U.S. Geological Survey Map created by Tyler Lewis/USGS. (d) Data (1995-2014) (Scanlon et al. 2007).

Empirical evidence for this hypothesis, however, is ambiguous (Maestre and Escudero 2009, Meloni et al. 2017*b*, Weerman et al. 2012). Additionally, simulation studies in more complex models suggest that details of systems matter, thus questioning the generality of these cluster based indicators (Génin et al. 2018*b*, Schneider and Kéfi 2016). Nevertheless, the possibility of inferring ecosystem resilience from a single snapshot and the increasing availability of low-cost remotely-sensed spatial datasets, where these methods can be applied, is attractive. Therefore, an evaluation of the generality and robustness of clustering properties as a signature of ecosystem resilience is needed.

To understand issues of generality, we must discuss another class of power-law behaviours that are considered universal features near/at *critical points* of phase transitions. Here, we emphasize that the theoretical underpinnings of ecosystem dynamics and indicators of stability are based on principles derived from the theory of phase transitions and bifurcations (Scheffer et al. 2009, Strogatz et al. 1994). This theory predicts that as a system nears a critical point of phase transitions, it takes increasingly longer to recover from perturbations. This phenomenon of slowed recovery is called *critical slowing down* (CSD) in the context of continuous phase transitions in the physics literature. However, a similar effect of slowed recovery appears even in ecological models that show abrupt transitions (Scheffer et al. 2009, Strogatz et al. 1994, Wissel 1984). Consequently, CSD has been widely used to devise methods to detect the approach of critical thresholds in ecosystems (Scheffer et al. 2009, Wissel 1984). An aspect of CSD that is much less known in the ecology literature is that close to, and at the critical point, the strength of perturbation decays as a power-law function of time-indicating a very slow recovery (Ma 2000, Sethna 2006, Stanley 1999); this is in contrast to systems far away from thresholds where perturbations decay exponentially fast. In fact, many power-law behaviours arise near/at continuous phase transitions (Ma 2000, Sethna 2006, Stanley 1999).

We highlight an interesting contrast between the two power-law relationships we have discussed thus far: While the power-laws associated with CSD are expected to *emerge near/at critical points* of phase transitions (Ma 2000, Sethna 2006), the power-laws in clustering are hypothesised to be *lost near/at critical thresholds* of ecosystem collapse (Kéfi et al. 2007). It is now fairly well established that many mechanisms cause emergence of power-laws even away from critical thresholds (Newman 2005, Pascual and Guichard 2005, Roy et al. 2003). However, the theoretical basis for why a loss of power-law clustering can indicate approach to a critical threshold in ecosystem models is unclear. Furthermore, elucidating relationships (if any) between the dynamical phenomenon of CSD and cluster size properties, has not gained attention in the literature. Such an exercise will not only prove helpful in evaluating the generality of ecosystem resilience indicators but also reveal the crucial role of local positive feedback in ecosystem patterning.

In this article, we review and synthesize the literature on how local ecological processes lead to the formation and dynamics of clusters, and how the resulting spatial patterns relate to ecosystem stability. Owing to the interdisciplinary nature of the study, we introduce important terms and concepts via summaries in Boxes 1, 2 and 3 and a glossary in Table 1.

We summarize our main findings here: First, our synthesis reveals the importance of local positive feedback in the emergence of power-law clustering in various ecosystems. To probe the relationship between positive feedback, clustering and resilience, we use a spatially-explicit model which, unlike previous relatively complex models, decouples the effects of positive feedback and environmental stress. Together with synthesis of previous studies, our analyses enables us to demonstrate that power-law clustering (or loss thereof) is unrelated to resilience. We then demonstrate how CSD - a universal feature of dynamical systems near thresholds - manifests as a power-law decay of spatial correlations. We discuss the important role of positive feedback in shaping clustering properties and suggest future directions of research to quantify patterns/dynamics of clustering and to infer ecological interactions.

### Glossary

1. Regime shifts: Changes in qualitative nature of ecosystem states. These changes can be abrupt or gradual functions of the underlying drivers.
2. Critical threshold: The value of an environmental condition (such as rainfall) and/or state variable (e.g. woody cover) at which a system undergoes an abrupt regime shift. In some ecology papers, it is used interchangeably with critical point but here we avoid doing so.
3. Resilience: The amount of change a system can withstand without transitioning to an alternative state. In the model described in Box 2, we interpret resilience as the distance to the threshold driver (or density).
4. Stability: The rate at which a system recovers to its original equilibrium from small perturbations.
5. Critical point: In the physics literature, this term refers to the value of driver at which the system typically undergoes a *continuous* phase transition from one state to the other.
6. Critical slowing down: The phenomenon in which systems near threshold of transitions are slow to recover from perturbations.
7. Positive feedback: Interactions between individuals that resuls in enhanced reproduction and/or reduced death rates of both individuals.
8. Cluster: A set of individuals who are within a minimum distance (typically the nearest neighbor distance) of at least one member of the same set.
9. Scale-free: A quantity having infinite average value, thus lacking a characteristic scale. Also see Box 1.
10. Percolation: In the physics literature, percolation is the movement/spread of an agent through the entire extent of the system via a connected path of sites.
11. Percolation density: The lowest density of occupied sites at which a fully connected path in the system is possible. At the same density, we observe a scale-free distribution of cluster sizes in the landscape.
12. Spatial autocovariance function: Covariance between states at two locations as a function of the distance between them. Also see Box 3.
13. Power spectrum/Spectral density function: Strength of fluctuations as a function of frequency; it is the Fourier transform of the autocovariance function. Also see Box 3.

#### BOX 1: POWER-LAW AND SCALE-FREE BEHAVIOURS

Biology is replete with examples of self-organised spatial clustering (Guichard et al. 2003, Rietkerk and van de Koppel 2008, von Hardenberg et al. 2001). In some cases, clumps have a wide range of sizes such that the frequency of occurrence of clumps of a particular size (denoted by *x*) decays as a power function of the size i.e. *f*(*x*) = *cx*^−*β*^ (defined for all clusters above a size *x* > *x*_*min*_ with *c* and *β* being constants). Below we describe two interesting properties of this function.

### Heavy-tailedness

The power-law frequency distribution has much higher occurrences of extreme events than predicted by commonly used distributions such as Gaussian or exponential distributions (Fig 2); this feature of the power-law distribution is also called *heavy-tailedness*.

**Figure 2:**
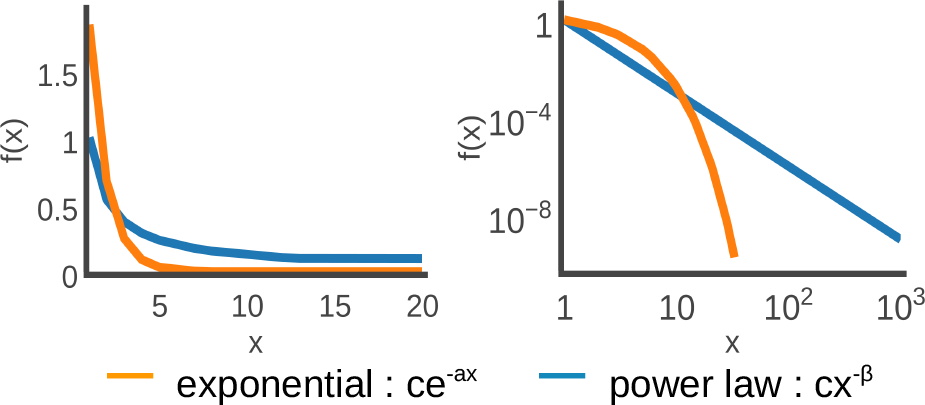
The plot on the left shows that the power-law function has a heavier tail, i.e. higher frequency (*f*(*x*)) of occurrence of large events, than in an exponential function. The plot on the right shows that power-law function is a straight line on log-log axes; the heavier tail of power-law is evident here too.

### Scale-free power-laws

Power-laws with an exponent *β* ≤ 2 mathematically describe features *that lack a characteristic size/length scale*. To see this, we observe that when *β* ≤ 2 the mean of this distribution is infinite. Exact expressions for the mean (*x̄*) and variance 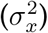 of the (normalised) power-law probability density function, denoted by *p*(*x*), are given by

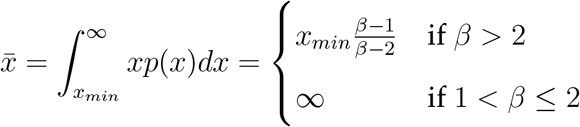

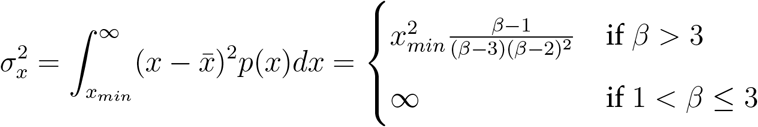

Thus, there is no characteristic size or typical length scale in this distribution, when *β* ≤ 2, and therefore the distribution is called *scale-free*. Power-law distributions of biological quantities with exponents *β* ≤ 2 are therefore intriguing. Such distributions, however, are not uncommon and have been documented in various ecosystems (Fig 1).

#### BOX 2: SPATIAL MODEL WITH POSITIVE FEEDBACK

Several spatial models in ecology try to explain power-law clustering but due to their relative complexity, it is difficult to clearly elucidate the role of positive feedback on clustering and resilience (Guichard et al. 2003, Kéfi et al. 2007, Manrubia and Solé 1997, Scanlon et al. 2007). To address this problem, we employ a simple spatially-explicit model with only two parameters. In this model, we consider a discrete two-dimensional space where each grid cell is updated probabilistically depending on states of cells in its neighbourhood. The simplicity of this model allows us to independently tune, and thus study effects of, environmental driver and positive feedback on spatial patterns via two parameters *p* and *q*, respectively. See Fig 3 for a schematic of the update rules; detailed model description is available in Appendix A and was first described in the physics literature in Lübeck (2006) and has been recently adopted in the context of regime shifts (Eby et al. 2017). Using this model, we study the effect of positive feedback (*q*) on steady-state density (defined as proportion of occupied sites) and spatial patterns (quantified via cluster size distributions and spatial power-spectrum) as a function of the environmental driver *p*. We add that reducing *p* in this model can also be interpreted as increasing environmental stress.

#### Positive feedback and abrupt regime shifts

Stronger positive feedback in ecosystems are known to cause non-linear and even abrupt responses to stress (Kéfi et al. 2010, 2016, Xu et al. 2015*b*). In our model too, when positive feedback is weak, the system undergoes a continuous transition from an occupied to a bare state as we increase environmental stress (Fig 4a). As positive feedback strength increases, the system can maintain a high density state even for higher levels of stress; but the system also exhibits an abrupt transition to a bare state when the stressor crosses the critical threshold. Henceforth, we refer to the point of transition (defined by either driver value (*p*) or density (*ρ*)) from an occupied to a bare state as *threshold*. When we specifically refer to a continuous transition, we call it a *critical point* whereas the corresponding term for the discontinuous transition is *critical threshold* (also see Glossary).

**Figure 3:**
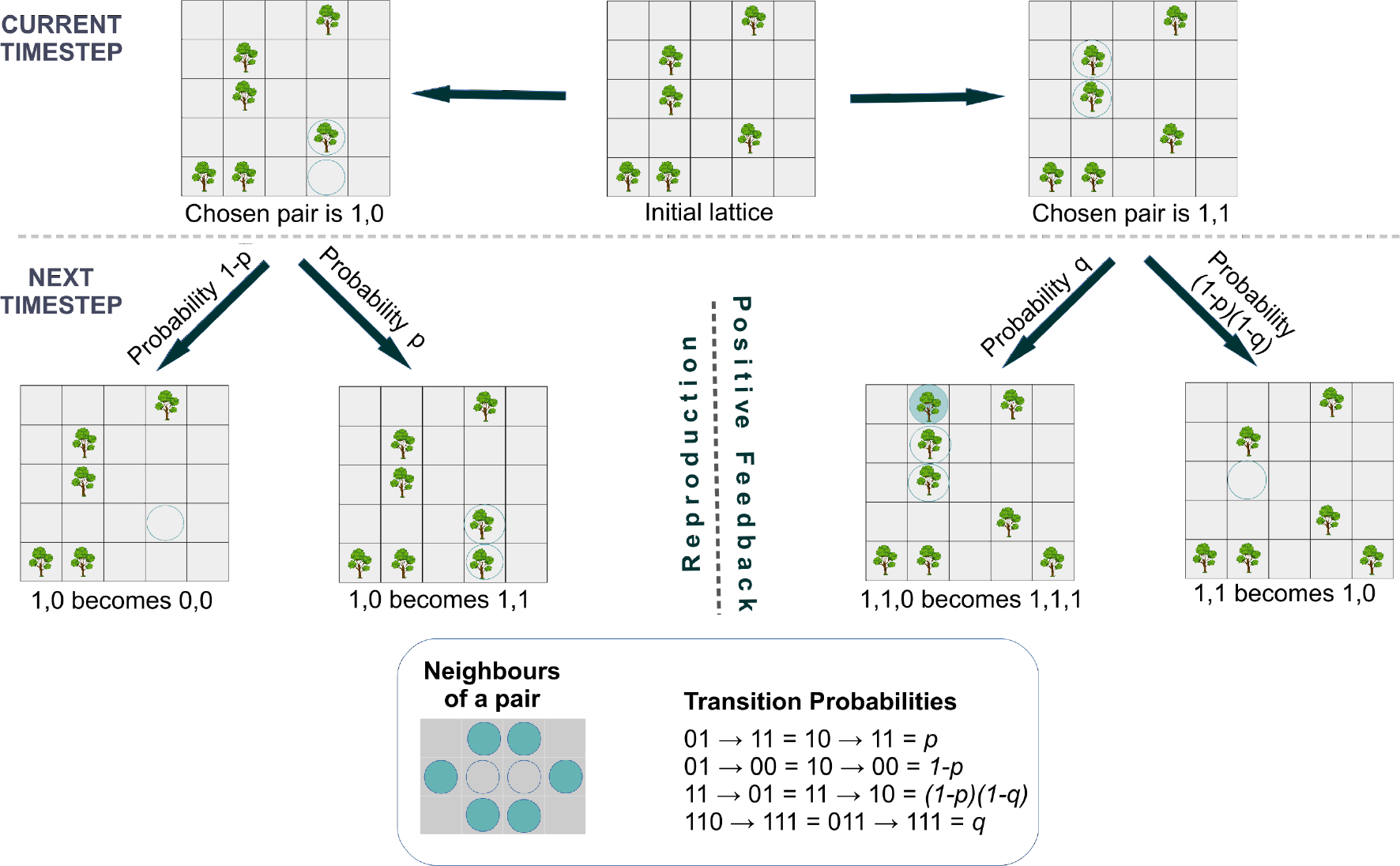
Schematic representation of the model and simulation procedure, for a given ‘Initial lattice’ shown at the centre of the top row. The parameter *p* represents baseline birth rate whereas *q* represents the strength of local positive feedback; reducing *p* in this model can be interpreted as increasing environmental stress. Light blue circles represent (randomly) chosen cells to update. Depending on the states of chosen cells, the update scheme results in baseline birth or death (left part of second row), or increased birth or reduced death due to positive feedback (right part of the second row). The box at the bottom shows (i) neighbours of a focal pair of cells and (ii) model udpate rules captured via transition probabilities.

**Figure 4:**
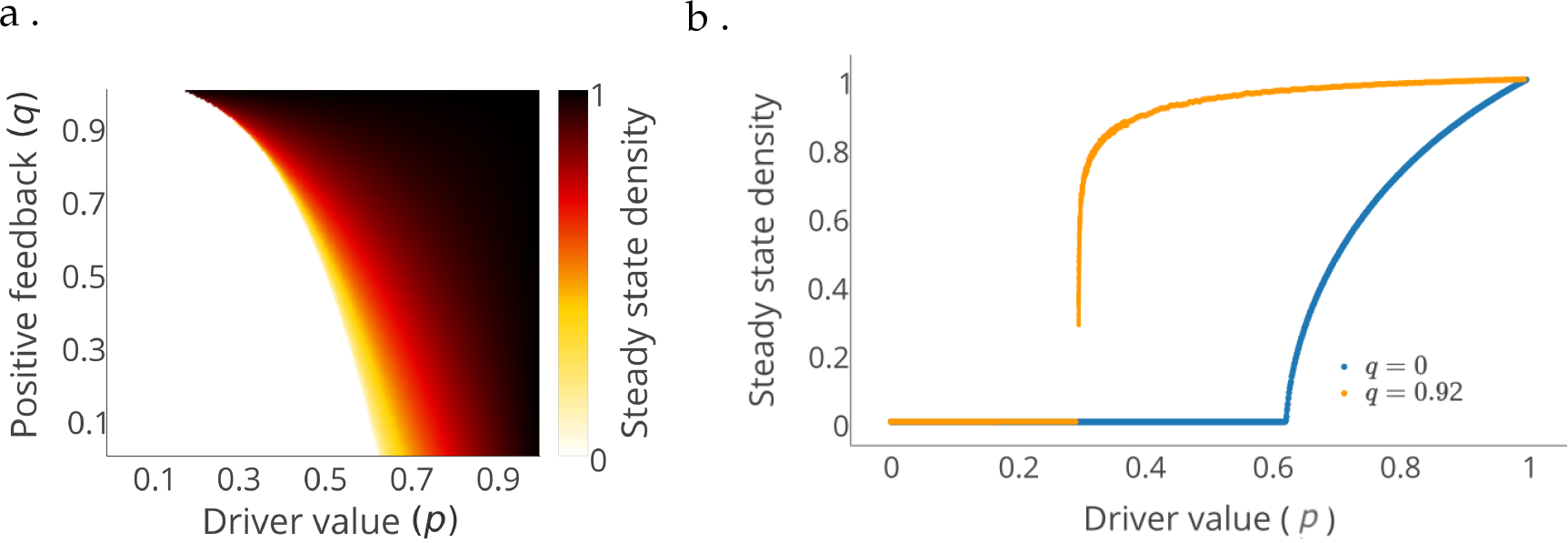
Positive feedback increases the non-linearity and cause an abrupt collapse in response to stress. (a) Steady-state density as a function of driver value (*p*) and positive feedback (*q*). (b) shows a closer look for two values of positive feedback, *q* = 0 (continuous transition) and 0.92 (discontinuous transition). Lattice size 1024 × 1024.

## III. POSITIVE FEEDBACK PROMOTES POWER-LAW CLUSTERING AT RELATIVELY LOW DENSITIES

Power-law cluster size distributions are seen in diverse ecosystems, including drylands, mussel beds, seagrass beds, sawgrass and forest fires (Fig 1). Many studies propose that these systems are likely shaped by positive feedback (Aguiar and Sala 1994, Boada et al. 2017, Dell et al. 2016, Foti et al. 2013, Guichard et al. 2003, Kéfi et al. 2007, Knowlton 2004, Maestre et al. 2003, Scanlon et al. 2007). For example, in semi-arid landscapes, seed germination and seedling survival probabilities are higher in the neighbourhood of other plants than out in the open (Aguiar and Sala 1994, Maestre et al. 2003). This results from reduced light and heat stress as well as increased water availability to young saplings in the vicinity of adult plants. Similarly, in mussel-beds, steadfast attachment of mussel to the substrate is directly dependent on the attachment of neighbours (Guichard et al. 2003). In macroalgal beds, recruitment and survival of macroalgal fronds shows density dependence due to the protection offered by neighbours from herbivory by sea urchin and fish (Boada et al. 2017, Dell et al. 2016).

To understand how positive feedback promotes such clustering, it is insightful to first discuss how power-law cluster-size distributions are also realised in ‘null models’ that are devoid of any interactions among organisms. In spatial null models, individuals are initially assigned to random locations on a two-dimensional discrete lattice. They then either die or give birth to an offspring at a rate that does not depend on presence/absence of any individual on the landscape (Kéfi et al. 2011). Consequently, the proportion of occupied sites in the landscape (henceforth called density) changes from nonzero values to zero (a bare state) as a gradual function of decreasing birth (or increasing death) rates (Grimmett 1999, Kéfi et al. 2011). These null models correspond to classic models in the physics literature in the context of a phenomenon called percolation (Stauffer 1979). The lowest density at which there is a non-zero probability of emergence of a fully connected path in the system is called the *percolation density*; at the percolation density (denoted by *ρ*_*p*_), the system also shows a scale-free clustering. In other words, despite the lack of positive feedback in these null models, a powerlaw cluster size distribution with *β* < 2, and hence scale-free clustering, occurs at the percolation density (*ρ*_*p*_). The value of percolation density depends on the geometry of the landscape. For ecological contexts, a relevant geometry is that of two dimensional square lattice where the percolation density is 0.59 (Grimmett 1999, Stauffer 1979).

In many ecosystems, densities that correspond to power-law clustering are typically lower than the above mentioned percolation density of the null model. For example, regions in the Kalahari show power-law cluster-size distributions of vegetation for densities ranging from 0.14 to 0.54 (Scanlon et al. 2007); a bulk of these areas also exhibit power-laws with exponent *β* < 2, and are thus scale-free (see Box 1 and Glossary). Power-law cluster-size distributions observed in several other ecosystems also show exponents within the scale-free range (e.g. Fig 1). To explain such power-law clustering, many spatial ecological models of ecosystems have been developed (Grassberger 1993, Guichard et al. 2003, Kéfi et al. 2007, Manrubia and Solé 1997, Scanlon et al. 2007). These models often incorporate ecosystem-specific processes and are consequently complex, involving many parameters. Nevertheless, they have commonalities. For example, they all assume local positive feedback in some form that causes increased birth (or reduced death) rates of individuals who are surrounded by others (Box 2). Below, we explain how local positive feedback can lower the percolation density.

The emergence of power-law clustering depends on how local interactions between individuals scale to cluster dynamics. Even in spatial null models that are devoid of any positive interactions, clusters form entirely due to random filling of the lattice; furthermore, larger clusters are more likely to merge with other clusters and therefore have higher growth rates. Theory predicts that power-law clustering emerges whenever clusters grow in proportion to their size, a phenomenon known as proportionate growth (Grimmett 1999, Manor and Shnerb 2008, Stauffer 1979). Such growth occurs at the percolation density of 0.59 for the spatial null models for a square lattice. In models with local positive interactions, empty sites near an existing cluster of occupied states are more likely to become occupied. This not only expands the original cluster but also increases chances of merger of this cluster with a nearby cluster. This dynamic of clusters is contrast to the spatial null model where expansion as well as merger of clusters are driven entirely by the random filling of the landscape. Therefore, in models with positive feedback, proportionate growth and scale-free cluster size distributions (i.e. a power-law with 1 < *β* < 2) occur at densities lower than the percolation density of the null model (Scanlon et al. 2007). This may offer a potential explanation for the observed low densities at which power-law clustering is seen in many ecosystems (Fig 1).

We support this argument by showing how percolation density changes with positive feedback in our model (Box 2). To do so, we use the concept of *percolation probability* which is defined as the probability of occurrence of a fully-connected path of occupied cells in the landscape. In Fig 5, we display the percolation probability as a function of density for two different values of positive feedback and the spatial null model. We then identify percolation density as the lowest density at which this probability is non-zero. We find that the percolation density is lower for the system with higher positive feedback, consistent with our synthesis of previous theoretical and empirical studies discussed above. Interestingly, we also observe that weak positive feedback leads to continuous change in percolation probability whereas strong positive feedback, owing to stronger nonlinear response of the system, makes it discontinuous (Fig 5).

**Figure 5:**
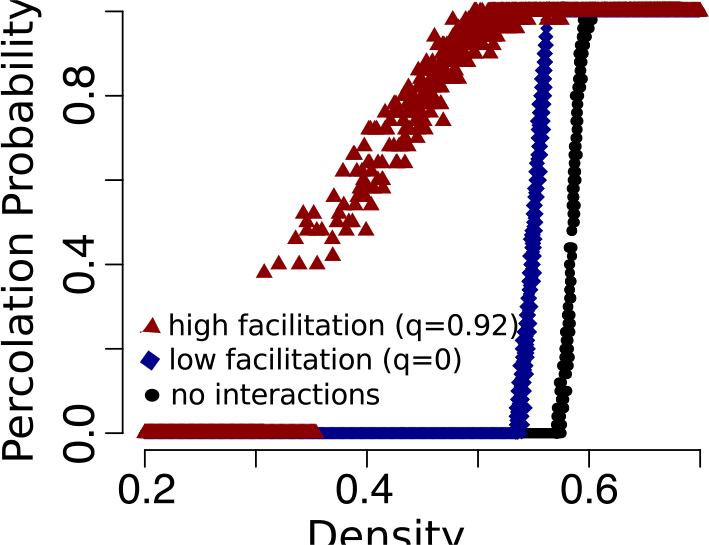
Percolation probability changes from zero to nonzero at density 0.59 for the spatial null model, 0.53 for low positive feedback (*q* = 0) and 0.31 for high positive feedback (*q* = 0.92). For each of these cases, these transitions in percolation probability occur exactly at the density where power-law cluster size distributions are observed; see Fig 6. Lattice size 256 × 256.

Putting empirical studies together with spatially-explicit models of clustering, we conjecture that strong positive feedback is likely to be the key interaction lowering percolation density in many ecosystems.

## IV. CLUSTER-SIZE DISTRIBUTIONS ARE NOT INDICATORS OF ECOSYSTEM RESILIENCE

Let us now use the above link we established between positive feedback and cluster-size distributions to address the larger question: how general is the relationship between cluster-size distributions and ecosystem resilience?

Studies over the last decade have suggested that fragmentation of large clusters leads to a thinning of the tail of the cluster-size distribution. Consequently, this causes loss of a power-law clustering, which can be used as an indicator of a stressed and less resilient ecosystem (Génin et al. 2018*a*, Kéfi et al. 2014, Kéfi et al. 2007). A corollary to this hypothesis is that ecosystems with power-law clustering are relatively farther from critical thresholds, and hence are likely to be more resilient. The evidence for this hypothesis in both models and data, however, has been ambiguous (Maestre and Escudero 2009, Meloni et al. 2017*b*, Moreno_de las Heras et al. 2011, Schneider and Kéfi 2016). Therefore, the generality of the relationship between cluster sizes and resilience remains unknown.

To resolve this, let us first consider how positive feedback affects both spatial clusters and resilience. As we argued in section III, positive feedback lowers the percolation density. Additionally, positive feedback promotes abrupt transitions and increases the threshold value of density from which the regime shift happens (Fig 4). Therefore, we hypothesize that, depending on the strength of positive feedback, power-law clustering can occur at any distance from the threshold of regime shift. We make predictions for two scenarios: we predict that in systems with *weak positive feedback*, the distance between percolation density and threshold of regime shift will be relatively large. Thus, increasing stress and an approach to threshold follows the previously expected pattern of loss of power-law clustering (Kéfi et al. 2014). In contrast, for systems with *strong positive feedback*, which are most likely to exhibit abrupt transitions, the distance between percolation density and the critical threshold of collapse will be negligble or even zero. Hence, power-law clustering may occur at the critical threshold itself and the loss of power-law clustering cannot be used as a resilience indicator.

To buttress our arguments, we analyse the model presented in Box 2. Indeed, our model analysis confirms our expectations: A weak positive feedback scenario shows that percolation density (*ρ*_*p*_) is relatively far from the threshold of transition (*ρ*_*p*_) (Fig 6a and inset); moreover, we find that loss of power-law clustering and appearance of thin-tailed (exponential) cluster-size distribution precedes the transition (see Appendix C). Our model reveals that this distance between the density of threshold of transition and percolation density reduces as a function of positive feedback and becomes even zero for large values of positive feedback (Fig 6c, d). Consequently, the qualitative features of cluster size distribution (e.g. being a power-law, truncated power-law or exponential) do not follow a general trend as a function of ecosystem stress see Appendix C. In Fig 6b and inset, we show a case where a strong positive feedback scenario shows a power-law clustering occurring very near, even possibly at, the critical threshold of collapse. Put together, our model analyses suggests that the relationship between cluster-sizes and ecosystem resilience heavily depends on the strength of positive feedback in the ecosystem. We recall that systems with strong positive feedback are most likely to exhibit abrupt shifts; it is precisely in these systems that the expected trend of cluster-size distributions, of loss of power-law as the system approaches thresholds, is least likely to be true. This questions the generality as well as potential utility of cluster-size distributions as indicators of ecosystem resilience.

**Figure 6:**
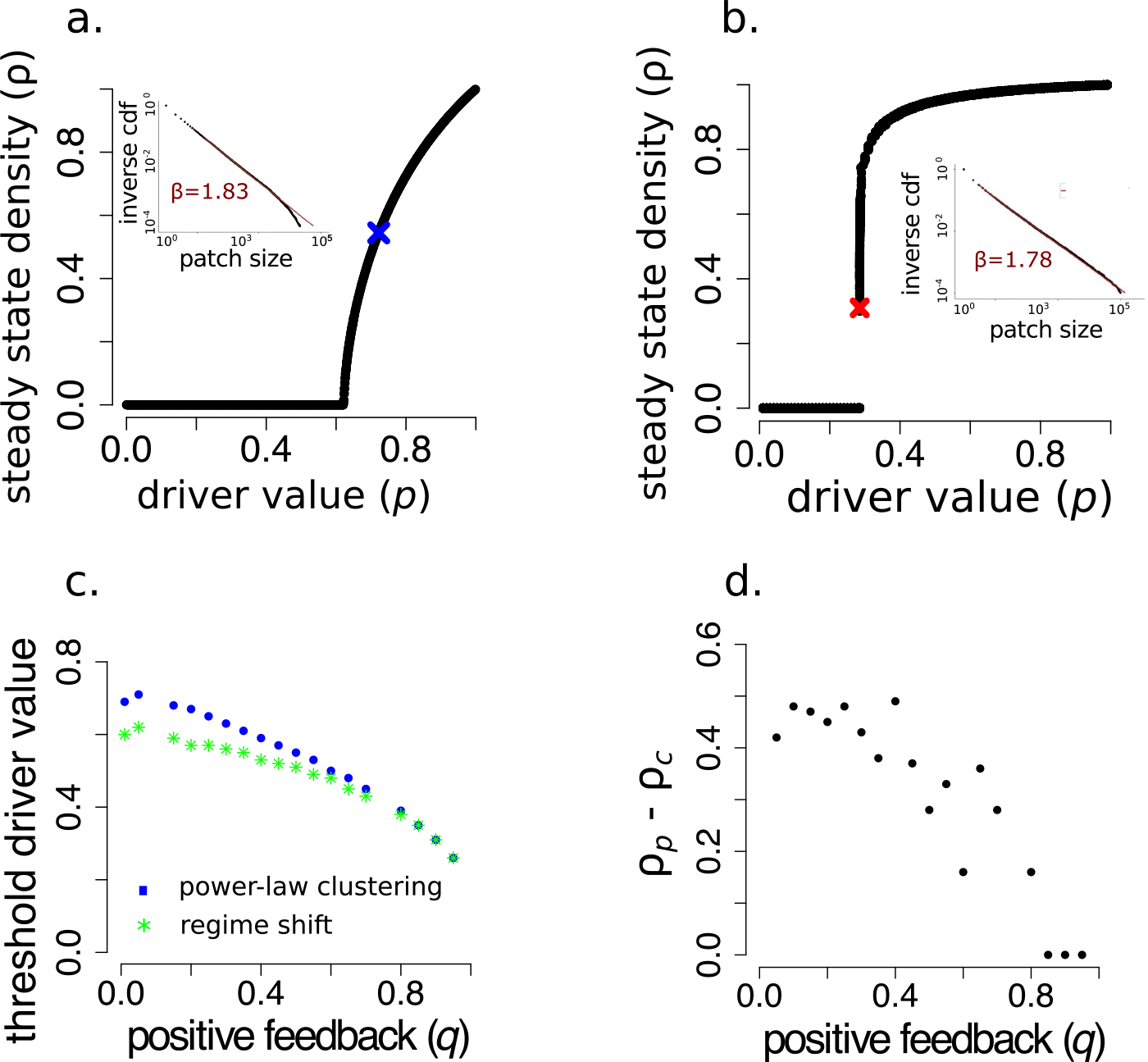
Strength of positive feedback, rather than distance to thresholds, determines the density at which power-law (scale-free) clustering occurs. The values of driver and density at which we find a power-law distribution are shown as crosses in the phase-diagrams (a) and (b), with their insets showing the corresponding inverse cumulative distribution function (CDF) of the patch-sizes. (a) When positive feedback is weak (*q* = 0), power-law clustering occurs far from ecosystem transition, consistent with previous hypotheses. (b) When positive feedback is strong (*q* = 0.92), power-law clustering can occur close to (or even at) the critical threshold of collapse. For the fitted function *kx*^−*β*^ wherein 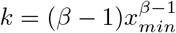, *x*_*min*_ = 17 in (a) and 3 in (b); lattice size used: = 1024 × 1024. (c) shows the driver values at which power-law clustering moves closer to the threshold of transition as positive-feedback (*q*) increases. (d) shows that *ρ*_*p*_ − *ρ*_*c*_, i.e. the difference between the density at which patches follow a power-law distribution (*ρ*_*p*_) and the density of the transition threshold (*ρ*_*c*_), reduces as positive feedback (*q*) increases. For (c) and (d), lattice size of 256 × 256 was chosen to reduce computational time. See Appendix C for cluster size distributions at other values of *p* and *q*.

We discuss above results in light of theoretical studies which too have found the association of cluster-size distributions with resilience to be tenuous (Génin et al. 2018*b*, Schneider and Kéfi 2016). These studies investigate spatially-explicit models of dryland vegetation and forest gap dynamics. They include, for example, a lowered grazing-induced mortality for individuals with more neighbours, a process termed associative protection. When the associative protection is high, they find power-law clustering at/near the critical thresholds of collapse. These results are consistent with our synthesis because associative protection in their model (i.e., reduced mortality for plants with neighboring plants) is analogous to increased positive feedback in our model (which causes reduced death rates for individuals with neighbors).

Synthesizing our results together with these recent studies, we argue that cluster-size distribution primarily depends on the strength of the positive feedback and that it cannot be employed as an indicator of ecosystem resilience. Furthermore, since cluster-size distributions do not primarily depend on proximity to critical threshold in these stochastic and spatial ecological models, we conclude that it is also unrelated to critical slowing down (CSD); we recall that CSD is a generic dynamical feature of systems near critical thresholds. See next section on how CSD influences spatial properties and causes power-law features in them.

## V. SCALE-FREE SPATIAL CORRELATIONS MAY ARISE AT CRITICAL THRESHOLDS OF ECOSYSTEM COLLAPSE

So far, we have demonstrated that cluster-size distribution do not represent resilience and hence cannot reliably indicate imminent regime shifts. However, the theory of phase transitions posits the emergence of scale-free features near/at critical points. Here, using our spatially-explicit ecological model, we illustrate how critical slowing down - a canonical features of dynamical systems near thresholds - causes scale-free behaviour in the spatial autocovariance function (Fig 7; Box 3;).

**Figure 7:**
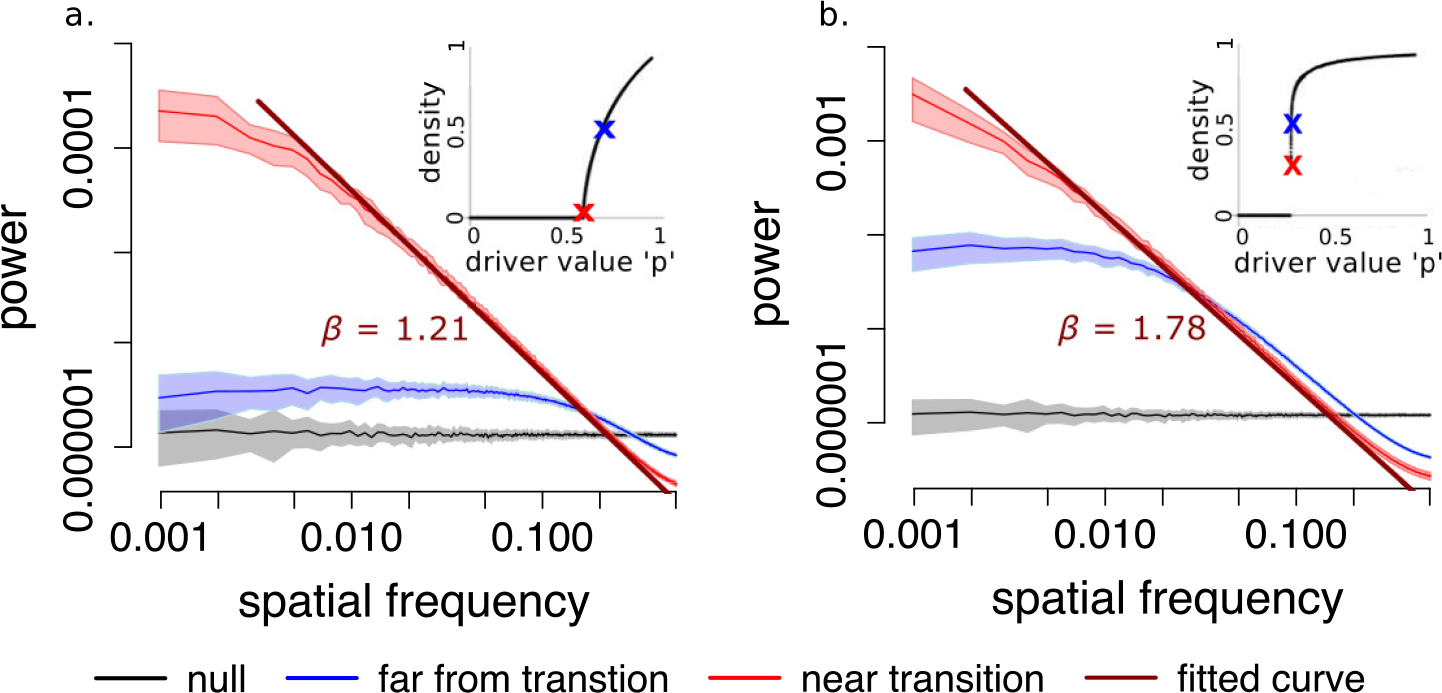
The power-spectrum of systems at very near (or at) the threshold of transitions decays as a power-law function of spatial frequency, (a) for *q* = 0 and (b) for *q* = 0.92. Lines represent the mean trend and bands, the SD. Insets show the location of parameter values for which power-spectrum are plotted. Blue is far from transition, Red is close/at the threshold and Grey represents the spatial null model. For the fitted function *kx*^−*β*^, *k* = 2.02 ∗ 10^−7^ when *q* = 0 and 4.65 ∗ 10^−8^ when *q* = 0.92. We used = 1024 × 1014 lattice.

As an ecosystem approaches a critical threshold, its return to equilibrium state, when perturbed, becomes increasingly slower. This phenomenon of critical slowing down (Ma 2000, Scheffer et al. 2009, Wissel 1984) has two implications - increased spatial correlations (Dakos et al. 2010) and increased spatial variance (Guttal and Jayaprakash 2009). To understand this, consider how a perturbation from the equilibrium state at any location in the ecosystem will spread in space. First, owing to slowed dynamics, the perturbation lives longer and, via spatial connectedness in the system, propagates to larger distances in the system (Ma 2000, Sethna 2006). Consequently, a measure of spread of perturbation, the correlation length, increases (Dakos et al. 2010, Ma 2000, Sethna 2006). Second, as the perturbations persist for longer duration, further disturbances enhance amplitudes of the fluctuations. This manifests as increasing spatial variance in the ecosystem as it moves towards the threshold (Guttal and Jayaprakash 2009). Here, we consider the spatial autocovariance function, defined as covariance of local densities at two locations separated by a distance *r* (Box 3). This function captures both spatial variance and correlations.

Before we illustrate computations of spatial autocovariance for our model, we make a couple of technical remarks. First, physicists often refer to the autocovariance function as the ‘correlation function’; some theoretical papers in ecology also do the same (Roy et al. 2003). Here, we have adopted the standard terminology that is used in quantitative ecology literature (Eq 2 in Box 3). Second, the theory of phase transition predicts critical slowing down and consequent scale-free behaviour at critical points of *continuous phase transitions* (Ma 2000, Sethna 2006). However, it has been shown that signatures of CSD are present, albeit with a relatively less magnitude, even in ecological models exhibiting abrupt transitions (Dakos et al. 2011, Scheffer et al. 2009). Consequently, we argue and demonstrate using the simple ecological model presented in Box 2 that scale-free behaviour may characterize critical thresholds of abrupt transitions as well.

Calculation of the spatial autocovariance function is often beset with statistical and computational difficulties. Therefore, we focus on a mathematically equivalent measure of correlations in spatial patterns via its power spectrum (Kéfi et al. 2014) (Box 3; Appendix D). It can be shown that the power spectrum is the Fourier transform of the autocovariance function (Reif 2009). The power spectrum of a spatial pattern provides a measure of the relative contribution of fluctuations at different spatial frequencies in the system, to its overall pattern. It is known in the ecology literature that as systems approach critical thresholds, the low frequency modes begin to dominate their power spectrum (Carpenter and Brock 2010, Kéfi et al. 2014). However, the full functional form of the power-spectrum is rarely quantified (but see Barbier et al. (2006), Bonachela et al. (2015), Couteron (2002) in the context of periodic and multi-scale patterns of dryland vegetation). Simulations of our model shows that the power-spectrum indeed becomes scale-free at critical thresholds for systems with both weak and strong positive feedback (Fig 7). We explain in Box 3 that a scale-free power spectrum is indicative of a scale-free autocovariance function. Thus, scale-free power-spectrum characterizes the structure of spatial perturbations near/at critical thresholds of ecosystem

### BOX 3: COVARIANCE, CORRELATION AND SPECTRAL FUNCTION

One way to capture the spread of disturbance in a system or the length scale of spatial fluctuations, is by constructing the spatial covariance function. The *spatial autocovariance function* for local density *ρ* for a distance *r* is defined as

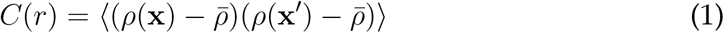

where 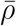 represents mean density over the entire landscape, angular brackets denote average over all locations x and x′ in the landscape that are separated by a distance *r*. Ecologists widely use the correlation function which is defined as

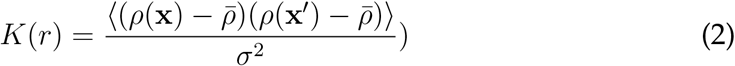

where *σ*^2^ is the spatial variance of densities in the ecosystem. Thus the covariance function is a production of the correlation function and the variance.

The *correlation length* is defined as the mean of the covariance function and can be interpreted as the average distance to which local fluctuations spread. The cor-relation length becomes infinite at the critical thresholds. This means that the covariance function then follows a power-law with an exponent less than 2 (Box 1).

The *power spectrum*, denoted by *S*(*k*), is the Fourier transform of its autocovariance function (Baugh and Murdin 2006, Reif 2009). Therefore, it can be calculated as

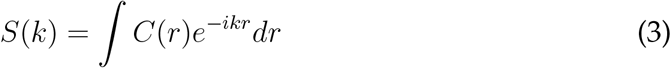

At critical thresholds, we expect the spatial covariance function to exhibit a power-law relation with distance

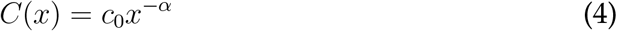

where *c*_0_ is a constant and *α* is an exponent less than two. The corresponding spectral function for an n-dimensional system is given by

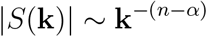

Therefore, evidence of a power-law spectral function is also evidence of a power-law autocovariance function.

## VI. DISCUSSION

In this study, we set out to investigate the generality of the conclusion that loss of power-law clustering in ecosystems is indicative of reducing resilience. First, our synthesis reveals that power-law clustering (or lack thereof) is unrelated to ecosystem resilience. We argue that this is because power-law clustering is fundamentally associated with local positive feedback rather than any generic dynamics of systems near critical thresholds of ecosystem collapse. Second, when ecosystems are in the vicinity of critical thresholds of collapse, critical slowing causes a power-law (scale-free) behaviour but in a different metric - the spatial autocovariance, or spectral function, of local densities.

### A. Local positive feedback, clustering and resilience

Previous ecological models that have attempted to resolve these connections include complex interactions often specific to particular ecosystems (Kéfi et al. 2011, 2007, Meloni et al. 2017*b*, Scanlon et al. 2007, Schneider and Kéfi 2016). In such models, many parameters contribute to local positive feedback and environmental stress, thus making it difficult to disentangle causal links between local processes and macroscopic patterns. Here, we deliberately used a simple model with only two parameters representing environmental stress and local positive feedback. The simplicity of the model we employed may also be seen as a limitation. However, it helped us conclude that loss of power-law clustering is not a robust indicator of approach to ecosystem transitions. Furthermore, it allowed us to disentangle the effects of environmental stress and positive feedback on clustering and resilience. Specifically, we propose a hypothesis that distance between power-law clustering (percolation threshold) and critical threshold of collapse reduces as the strength of positive feedback increases.

Seminal ecological models that try to explain power-law clustering observed in ecosystems (Kéfi et al. 2007, Scanlon et al. 2007) assume that local births/deaths of trees, in addition to being positively influenced by local density, is negatively regulated by *global-scale feedback*. Mechanisms such as rapid spread of water in the landscape (von Hardenberg et al. 2010) are offered as potential explanations for negative regulation of local growth due to global-scale vegetation density. Indeed, based on our synthesis ((Manor and Shnerb 2008, 2009) and Fig 6), we demonstrate that there is no need to invoke global-scale feedback; in fact, local positive feedback alone can explain the emergence of scale-free clustering in these systems.

Several empirical studies find neither scale-free clustering (Weerman et al. 2012, Xu et al. 2015*a*) nor the expected shifts of cluster-size distributions with increasing stress (Casey et al. 2016). However, they attribute this to an absence/disruption of global negative feedback in their systems (Casey et al. 2016, von Hardenberg et al. 2010, Weerman et al. 2012)(but also see Moreno-de las Heras et al. (2011)). As we argued in the previous paragraph, negative feed-back isn’t even a necessary condition for scale-free clustering. Further, based on our synthesis that cluster-sizes do not relate to resilience, we posit that these empirical results are not surprising.

Cluster-size distributions are nevertheless relevant in ecological contexts where connectivity or porosity of the landscape is of focal interest, such as in the case of forest fires or disease spread (Chaves et al. 2012, Turcotte and Malamud 2004). We illustrated that power-law clustering in our ecological model is associated with a transition in percolation probability, representing the emergence of a spanning cluster (i.e. a fully connected path) in the system. This is also seen in spatial models of predator-prey interactions (Roy et al. 2003). In the context of fire or disease outbreaks, presence of a cluster of vegetation (susceptible individuals) allows fire (disease) to easily spread within each cluster. Consequently, scale-free clustering, which indicates a highly connected landscape, allows the possibility of catastrophic fire (or disease) outbreaks. These models represent a fundamentally different class of models from what we have discussed in this paper (Dickman et al. 2000, Solé et al. 1999).

### B. Cluster-sizes and Critical slowing down

Our synthesis predicts that scale-free behaviour in spatial correlations, measured via autocovariance or spectral functions, can characterise critical thresholds. This feature, we argued, arises from the critical slowing down - i.e. slowed response of ecosystems near threshold points - which is a generic feature of many ecological transitions. How do scale-free correlations in density (described in V) and scale-free clustering (Section III) relate to each other? They both indicate emergence of large spatial scales in the system. However, they capture fundamentally different properties. Scale-free correlations in density indicate that *perturbations* spread to large distances in ecosystems as a consequence of critical slowing down. Therefore, it captures the dynamics of perturbations and hence can be used to infer stability or lack thereof. In contrast, scale-free clusters indicate the presence of large clusters, which do not correspond to dynamics of how perturbations decay. Therefore, clustering properties are unrelated to resilience, as we indeed demonstrate in Fig 6.

How efficient is it to use scale-free features of density correlations as an early warning signals (EWS) of regime shifts or critical transitions (Scheffer et al. 2009)? The purpose of early warning signals is to detect signatures of approach to critical thresholds. In that sense, computing simpler metrics of spatial autocorrelation between neighboring sites (Dakos et al. 2010) or spatial variance (Guttal and Jayaprakash 2009) may have advantages such as ease of computation and better statistical reliability in comparison to characterising the complete form of autocovariance or spectral functions. On the other hand, simpler metrics are also easily affected by external factors, such as increased spatial heterogeneity or external variability (Dakos et al. 2010, Kéfi et al. 2014) and hence confound interpretations. Further investigations can reveal the relative efficacy of different spatial metrics.

### C. Future directions

Our synthesis suggests some exciting directions for future research. The focus of recent research, as reviewed in this paper, has been to understand how local interactions produce clustering properties, and how clustering properties can be used to infer resilience. However, the inverse problem of inferring ecological interactions from spatial images of ecosystems remains poorly studied. For example, the occurrence of power-law or such heavy-tailed distributions can itself be used to infer the role of local facilitative interactions in the ecosystem. Indeed, one recent study does suggest that skewness of cluster-size distributions can suggest positive feedback in dryland-vegetation systems (Xu et al. 2015*a*). The bigger question remains open: can we quantify the strengths and spatial scales of positive feedback and other ecological interactions between organisms by analysis of spatial images, for example via geometrical properties of clusters such as cluster sizes, fractal dimensions of clusters, and the strength of spatial correlations in the system.

With recent advancements in remote sensing and reducing costs of spatial images, we can also procure extensive high-resolution spatial data over time. This will enable us to quantify not only patterns, as described above, but also dynamics of various cluster properties. Unlike static properties available from a single image, dynamical properties may reflect stability of ecosystems by capturing how systems respond to perturbations. Theoretical works, inspired by studies of domain growth in phase transitions, describe the dynamics of clusters in simple ecological models exhibiting continuous and discontinuous transitions (Manor and Shnerb 2008, Weissmann et al. 2017). However, much remains to be done in integrating these studies with real data. This requires extensive theoretical and computational studies to identify suitable metrics of clustering properties, development of statistical frameworks including appropriate spatial null models and finally, empirical validations/applications based on analyses of aerial images of ecosystems.

### D. Concluding remarks

Our synthesis helps us disentangle processes that generate power-law cluster sizes, scale-free correlations and how they relate to ecosystems’ critical thresholds. Real world analyses however can sometimes yield misleading patterns, including power-laws and scale-free behaviours due to sampling artefacts (Plank and Codling 2009) or misfitting (Clauset et al. 2009, Meloni et al. 2017*a*, White et al. 2008). Various ways in which patterns are misconstrued as power-laws have been discussed in detail in multiple other forums (Breed et al. 2015, Clauset et al. 2009, Stumpf and Porter 2012). Where there is a true power-law with an exponent less than two, since it is indicative of diverging quantities, there is a tendency to associate such a pattern with a critical phenomenon. However, scale-free patterns can also arise when underlying processes operate at multiple scales and due to landscape heterogeneity (Khaluf et al. 2017, Petrovskii et al. 2011). Naive association of observed scale-free behaviours with either criticality or stability is problematic. An additional challenge in interpreting spatial patterns in ecosystems is to disentangle effects of underlying spatial heterogeneity from true self-organisation. With the increasing availability of high-resolution spatial datas, from satellites to drone based imagery, of various ecosystems, spatial analyses are likely to be widely deployed in the future. Our study highlights the importance of having a clear understanding of how local-interactions drive macroscopic behaviours to infer ecological interactions and resilience of ecosystems.

## VII. DATA AND CODES

All simulation analyses codes, with simulation datasets corresponding to results presented in this paper, have been made publicly available at: https://github.com/ssumithra/PowerLawCriticalityPaper. Detailed instruction on execution of these codes are also provided.

## VIII. ONLINE SUPPLEMENTARY MATERIALS

**Appendix A:** Power-law Vs exponential functions.

**Appendix B:** Detailed model description.

**Appendix C:** Statistical fitting of cluster-size distributions.

**Appendix D:** Cluster-size distributions across the phase-diagram for low and strong positive feedback.

**Appendix E:** Power-spectrum fitting.

## IX. ACKNOWLEDGEMENTS AND AUTHOR CONTRIBUTIONS

VG acknowledges research support from a DBT-Ramalingaswamy Fellowship, DBT-IISc partnership program, ISRO-IISc Space Technology Cell and infrastructure support from DST-FIST. SS and SM were supported by a scholarship from MHRD via IISc. *Author Contributions*: SS conceived the idea. SS wrote codes with key contributions from SM (model), AV (statistical fitting) and VG (power spectrum). SS conducted analyses and produced figures. SS and VG synthesised the literature review and model results. SS and VG wrote the manuscript with comments from SM and AV. All authors gave final approval for publication and have no conflict of interests to declare. We thank Sonia Kefi for many insightful discussions. We thank Hari Sridhar and other lab members for comments on the manuscript.

