## Appendix for "Clustering and correlations: Inferring resilience from spatial patterns in ecosystems"

### SUPPLEMENTARY MATERIAL

Sumithra Sankaran,[1,*] Sabiha Majumder,[2]
Ashwin Viswanathan,[3] and Vishwesha Guttal[1]

[1]*Centre for Ecological Sciences, Indian Insititute of Science, Bangalore, India*

[2]*Department of Physics, Indian Insititute of Science, Bangalore, India*

[3]*Department of Environmental Systems Science, ETH Zurich, Switzerland*

#### CONTENTS

A. Detailed model description 2

B. Statistical fitting of cluster-size distributions 3

C. Cluster-size distributions 5

D. Power-spectrum fitting 6

1. Resilient systems (far from transition points) 6
2. Critical systems (very near/at transition points) 7
3. Comparing Lorenztian and scale-free spectra 9

References 10

*

#### Appendix A: Detailed model description

The system is modelled as a two dimensional discrete space consisting of $N$ x $N$ cells. Each cell can occur in any of two possible states: unoccupied (or density $0$) and occupied (or density $1$). The cells are updated probabilistically and the simulation sequence runs as described below.

**Step 1**: Select a cell at random.

**Step 2**: If this cell is unoccupied, return to step 1. If occupied, proceed to step 3.

**Step 3**: Select at random, one of the four nearest neighbors of the chosen cell cell.

**Step 4**: If this selected neighbor cell is empty ($0$), update its state to $1$ by probability $p$, else update the state of the first chosen cell to $0$ (the probability of this is then $1 - p$). Return to step 1 for a new iteration.

**Step 5**: If the first chosen cell (in step 1) and the neighbor cell (selected in step 2) are both occupied, (a) with probability 'q' select one of the six possible nearest neighbors of the pair and make it occupied (if it it was already occupied, nothing changes), else update the first chosen cell (in step 1), to $0$

The cells are, thus, updated asynchronously. Each discrete time step implies that the update rules are iterated $N^2$ number of times sequentially; this procedure ensures that, on an average, all sites are updated once per discrete time step.

We ran our simulations with a $1024 \times 1024$ system size at increasing resolutions of $p$ and $q$ to construct the steady-state phase diagrams of mean density of the landscape and identify the critical driver values. The system was considered to have reached a steady-state once the mean density had saturated. Cluster-size distributions and Power-Spectrum functions were computed from 50 replicate spatial snapshots of the system at steady-state. To compute the spanning cluster probabilities alone, we ran another set of simulations with $256 \times 256$ system-size and calculated these probabilities from 25 replicate spatial snapshots of the system at steady-state.

##### Appendix B: Statistical fitting of cluster-size distributions

Cluster size distributions were fit using methods proposed in [3]. It is worth recalling the methods of fitting power-laws have attracted much scrutiny in the literature. Therefore, we chose the statistical methods proposed by [3] which are widely accepted as rigorous to fit power-law distributions and to compare with other model candidates.

**Is power-law a good fit?**: First step in the process is to find out if power-law is even a good fit. The exponent of the distribution was estimated using Maximum Likelihood Estimation (MLE), and $x_{min}$ was identified by minimising the Kolmogorov–Smirnov(KS) distance between the fitted model and data. (The KS distance is the largest distance between the empirical distribution function of the data and the cumulative distribution function of the model). We assessed goodness of fit for our power-law model by re-fitting synthetic power law distributions (which we generated) with the same estimated exponent and xmin values. The fraction of synthetic datasets that result in a fitted model with a KS distance larger than the KS distance calculated when fitting our dataset, was considered the p-value of our fit. As described in [3], a p-value above 0.1 represents a good fit, and only when this condition was satisfied, we proceeded to compare with alternative models of cluster size distributions. Indeed, we found p-values of 0.49 for the low positive feedback model ($q = 0$)[see Fig 5a in main text] and 0.53 for the high positive feedback model ($q = 0.92$)[see Fig. 5b in main text]. This suggests good power-law fit of cluster size distributions for both values of positive-feedback, but one of them is away from critical point (Fig 5a) and the other is right at the critical point (Fig 5b).

**Is power-law the best fit?**: We compared power-law (PL) fit of the cluster size distributions with three different model fits: exponential (EXP), log-normal (LN) and power-law with an exponential cut off (PLE). Each of the candidate models was fit using MLE. Since power-law is a nested model of power-law with a cut-off, these two were compared using log-likelihood ratio. The other two candidate models were compared with the power-law model using Vuong test. We refer readers to [3] for further technical details. Visit our github page for reproducing our results: https://github.com/ssumithra/PowerLawCriticalityPaper

We know from percolation models that cluster size distributions typically shift from a bimodal to a power-law to a power-law with exponential cut off to an exponential as density in the system reduces [4, 5]. Thus, in our investigations of effects of positive feedbacks on clustering, these were obvious candidate distributions to fit to the data. In addition, we also fit and compare log-normal as a candidate function in order to make the reported results comparable with other studies discussed in [3]. However it must be noted that, till date, there exists no mechanistic processes that can yield a log-normal cluster size distribution in these systems . On the other hand, based on the theory of phase transitions, there is a well-reasoned expectation of scale-free behaviour at critical points.

Realising a true scale-free distribution requires, ideally, an infinitely large system. Our datasets consist of 50 replicates of identically sized lattices (of 1024 x 1024 cells). Even a true power-law distribution in these replicates would inevitably be best fit as a truncated power-law distribution, due to the limit imposed by the system-size. Given these finite size constraints in our data, a power-law as the best fit is inferred based on (a) the closeness of estimated parameters between the fitted power-law function and fitted power-law with exponential cut off function and (b) the range over which the power-law dominates the truncated power-law function.

For the data presented in Fig 5 of main text, power-law with exponential cut off was identified as the best fit model for both cases ($q = 0, p = 0.7225$ and $q = 0.92, p = 0.2852$). Given in Table 1 is the comparison statistics for the power-law (PL) fit with the other three considered models - exponential (EXP), Power-law with exponential truncation (PLE) and Log-normal (LN) fits.

| Dataset | PL Vs Exp<br>vuong test statistic | PL Vs PLE<br>log-likelihood ratio | PL Vs LN<br>vuong test statistic |
|---|---|---|---|
| **q=0, p=0.7225** | 34.31*** | -53.92 *** | -2.69 (p=0.996) |
| **q=0.92, p=0.2852** | 41.33608 *** | -9.71** | -15.22 (p=0.999) |

Table 1: Results of likelihood ratio test for fitted power-law vs other models. Positive values suggest that the power-law is a the better fit and negetive values favour the alternative model. Significance levels are as follows : '***' for $p < 0.001$, '**' for $p < 0.01$ '*' for $P < 0.1$ and '' ' for $P > 0.1$. For both datasets, log-normal could not be ruled out as a potential fit (p value given in brackets), while power-law with exponential cut-off was found to be the best fit

We see from the estimated rate (Table 2) of the fitted power-law with exponential cut-off model, $x^{-\beta}e^{-x/X_{max}}$ , that the $X_{max}$ is larger than the largest patch-size in the system. The power-law behaviour persists without any effect of the expo-

| Dataset | Fitted model | Exponent | Rate | $X_{min}$ |
|---|---|---|---|---|
| **q=0, p = 0.7225** | Power law with exponential cut off | 1.84 | $9.3 \times 10^{-6}$ | 17 |
| | Power law | 1.83 | | 17 |
| **q=0.92, p = 0.2852** | Power law with exponential cut off | 1.75 | $2.9 \times 10^{-6}$ | 3 |
| | Power-law | 1.77 | | 3 |

Table 2: Estimated parameter values of fitted power-law and power-law with exponential cut-off functions. The exponent estimates are very close for both model fits. The rate of exponential cut-off is very low, suggesting that the power-law persists over a large range of patch sizes.

nential truncation till atleast four orders of magnitude in patch sizes.

#### Appendix C: Cluster-size distributions

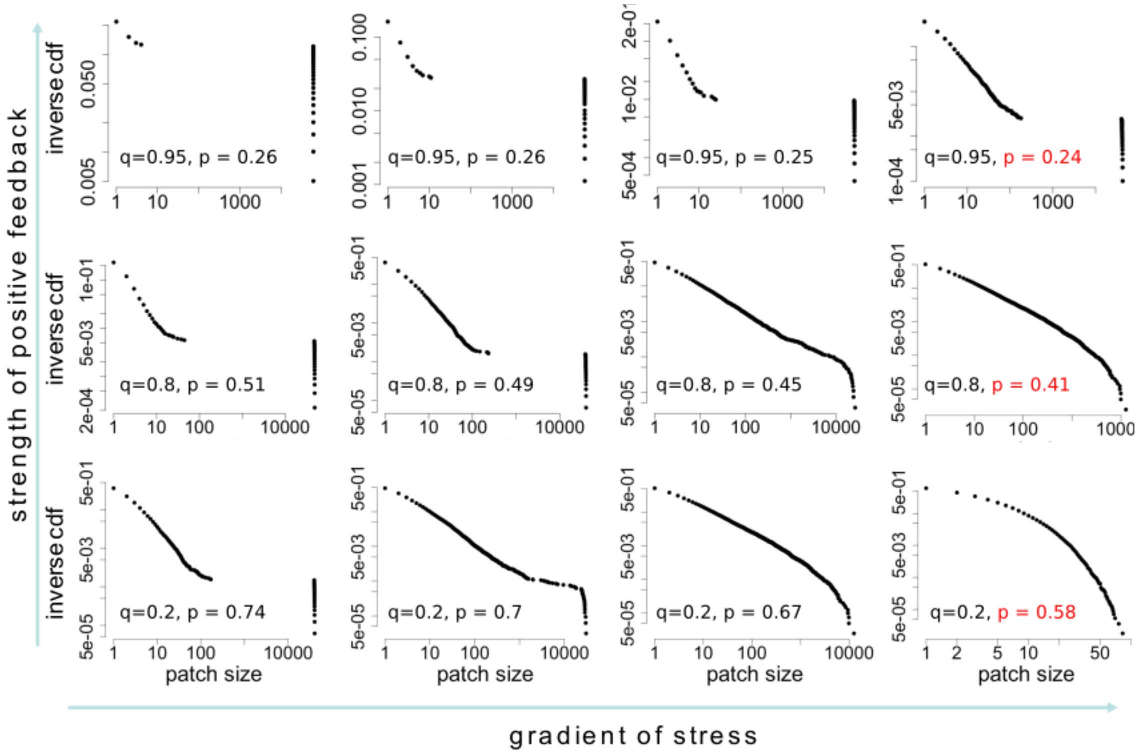


Figure 1: Plots show cluster size distributions realised in square lattices of 250x250 cells. The right most column of plots is of systems very near/at critical points, with the driver value shown in red. We see the entire range of cluster-size distributions: bimodal at very high density, far from the critical point to a power-law to a truncated power-law to an exponential, very near the critical point, when positive-feedback (‘*q*’ is low). As positive-feedback increases, 1. high densities prevail for lower and lower p values, 2. system begins to collapse from higher density states (see Fig 4 of main text). We then see power-law or, with very high positive-feedback, even bimodal clustering at the point of collapse.

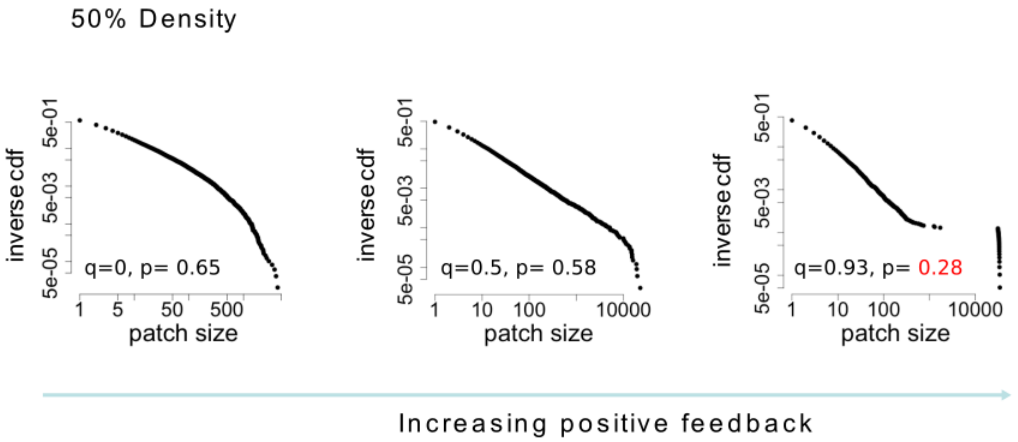


Figure 2: Plots show cluster size distributions realised in square lattices of 250x250 cells. At the same density, different cluster size distributions can be observed depending on positive-feedback strength (q). Since positive-feedback also results in the system collapsing from higher densities, increasing positive-feedback causes fat tailed distributions to occur closer and closer to the critical point.

#### Appendix D: Power-spectrum fitting

##### 1. Resilient systems (far from transition points)

It is well known that in systems far from transition the power-spectrum typically exhibits a Lorentzian functional form . Theoretically it is also expected that as the system reaches a critical point, its spectral function shifts to a power-law form. In our model too we found the Lorentzian function to be a good fit for resilient systems (both, with high and low positive feedback). The function was fit by running a non-linear least squared regression on the data. We present the results of the analyses below:

Dataset low positive feedback($q = 0$)
Formula: $y = k * a/((x - x0)^2 + a^2)$
Parameters:

| | Estimate | Std. Error | t value | Pr(>\|t\|) |
|---|---|---|---|---|
| k | 1.36x10$^{-6}$ | 1.6x10$^{-8}$ | 87.97 | <2e-16 *** |
| x0 | -0.2308 | 0.0051 | -45.35 | <2e-16 *** |
| a | 0.4116 | 0.0073 | 564.89 | <2e-16 *** |

Signif. codes: 0 '∗∗∗' 0.001 '∗∗' 0.01 '∗' 0.05 '.' 0.1 ' ' 1
Residual standard error: 0.0001061 on 25647 degrees of freedom
Number of iterations to convergence: 8
Achieved convergence tolerance: 0.0000000149

Dataset high positive feedback ($q = 0.92$)
Formula: $y\ k * a/((x - x0)^2 + a^2)$
Parameters:

| | Estimate | Std. Error | t value | Pr(>\|t\|) |
|---|---|---|---|---|
| k | $2.06\text{x}10^{-6}$ | $1.13\text{x}10^{-8}$ | 182.99 | <2e-16 *** |
| x0 | -0.0085 | 0.00017 | -49.67 | $2.72\text{x}10^{-11}$ *** |
| a | 0.0254 | $6.79\text{x}10^{-5}$ | 374.72 | <2e-16 *** |

Signif. codes: 0 '***' 0.001 '**' 0.01 '*' 0.05 '.' 0.1 ' ' 1
Residual standard error: 0.0004837 on 25647 degrees of freedom
Number of iterations to convergence: 8
Achieved convergence tolerance: 0.0000000149

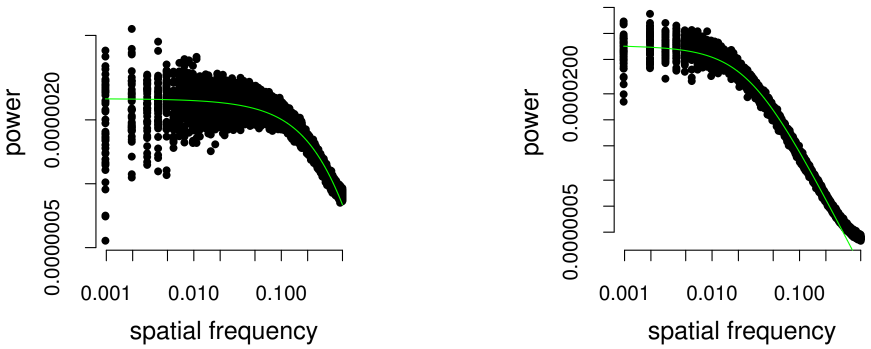


(a) $q = 0, p = 0.7225$ (b) $q = 0.92, p = 0.2865$

Figure 3: The lorentzian fit (green line) of power spectrum data (black dots) of resilient systems with low and high positive-feedback, see main text Fig 7. The plotted data correspond to the data shown in blue in Fig 7.

##### 2. Critical systems (very near/at transition points)

For systems near transition, as expected, we found that the power-law function was a good fit over a large range of the data. There are two approaches to fitting power-law relations: one is by log transforming the data and then fitting a linear model and the other, by directly fitting a non-linear model. Several studies demonstrate that log-transforming power-law distributions or frequency data violates assumptions underlying linear regressions such as normally distributed residuals. This, however, is not always the case. More precisely, fitting power-law data using non-linear least squares regressions assumes that the data have a constant standard deviation, whereas log transformation followed by linear regression is valid for data with a constant coefficent of variation [2]. Since our spectral data for systems near/at critical points, like other $1/f^{\beta}$ spectra [6], show an increasing trend of standard deviation with average power (see Fig.4), we used the linear regression method on log-transformed values. The range of the power-law function was estimated in the same way as described in the previous section, by identifying the x-value (spatial frequency) which minimises the KS distance between the predicted function and the data but an upper frequency limit of 0.3 was kept fixed for all datasets.

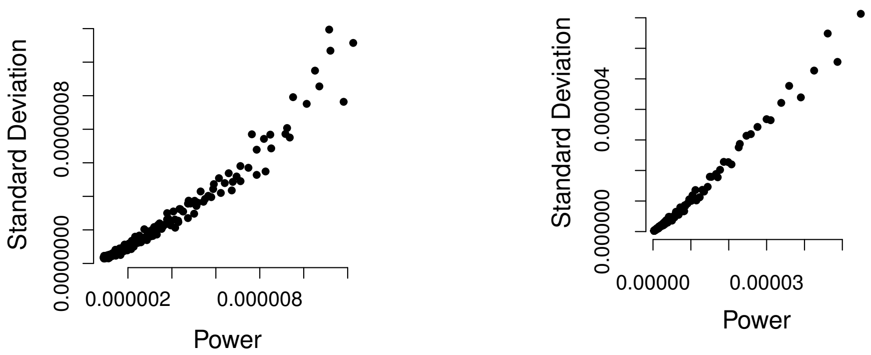


(a) $q = 0, p = 0.62275$

(b) $q = 0.92, p = 0.2852$

Figure 4: Varying $SD$ and constant coefficient of variation ($SD/mean$) of power-spectra of systems near transition.

Dataset: low positive feedback($q = 0$)
Formula: $y = (k * x^a)$
lm(formula = log(y)  log(x), weights = x)
Parameters:

| | Estimate | Std. Error | t value | Pr(>\|t\|) |
|---|---|---|---|---|
| $a$ | -1.2103 | 0.00096 | -1265 | <2e-16 *** |
| log $k$ | -15.4159 | 0.00159 | -9688 | <2e-16 *** |

Significance codes: 0 '***' 0.001 '**' 0.01 '*' 0.05 '.' 0.1 ' ' 1
Residual standard error: 0.01954 on 15698 degrees of freedom
Multiple R-squared: 0.9932, Adjusted R-squared: 0.9932
F-statistic: 1.6x$10^6$ on 1 and 15698 DF, p-value: < 2.2e-16

Dataset: high positive feedback ($q = 0.92$)
Formula: $y = (k * x^a)$
lm(formula = log(y)  log(x), weights = x)
Parameters:

| | Estimate | Std. Error | t value | Pr(>\|t\|) |
|---|---|---|---|---|
| $a$ | -1.7819 | 0.00128 | -1392 | <2e-16 *** |
| log $k$ | -16.8830 | 0.00283 | -5976 | <2e-16 *** |

Significance codes: 0 '***' 0.001 '**' 0.01 '*' 0.05 '.' 0.1 ' ' 1
Residual standard error: 0.1006 on 14798 degrees of freedom
Multiple R-squared: 0.9944, Adjusted R-squared: 0.9944
F-statistic: 1.938e+06 on 1 and 1 DF, p-value: < 2.2e-16

#### 3. Comparing Lorenztian and scale-free spectra

Comparison of the models (Lorentzian vs power-law) for the different datasets is impossible as the models not only have a different number of parameters, but are also not fit over the same range of data. One way to get around this is to consider the theory underlying the emergence of scale-free power-spectra in critical systems. Even data that follow a Lorentzian function, will follow a power-law over some (small) range of spatial frequencies. However, as the system approaches a critical point, low frequency interactions begin to dominate, thus increasing in power and leading to a shift in the spectrum such that the extent of the power-law region sharply increases (see section 4 on "scale-free spatial correlations in critical systems" in main text). Thus to compare the power-spectra behaviour in systems near/at the critical point with that of resilient systems, one can examine the range over which the power-law fit extends.

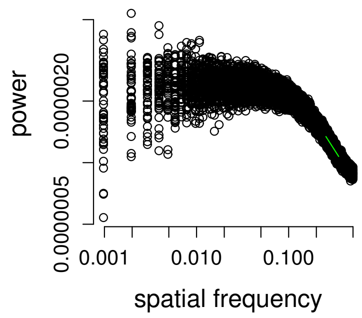


(a) $q = 0, p = 0.7225$, $PLR = 0.05$

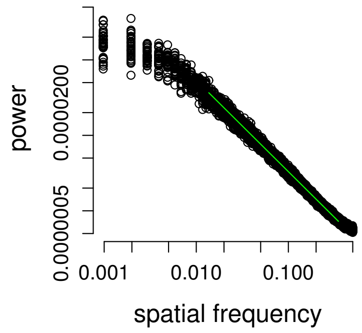


(b) $q = 0, p = 0.26675$, $PLR = 0.56$

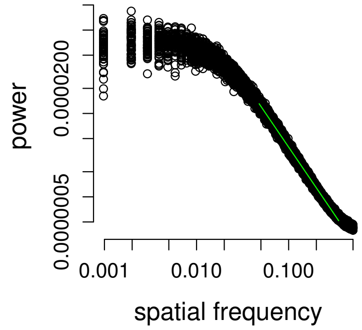


(c) $q = 0.92, p = 0.2865$, $PLR = 0.32$

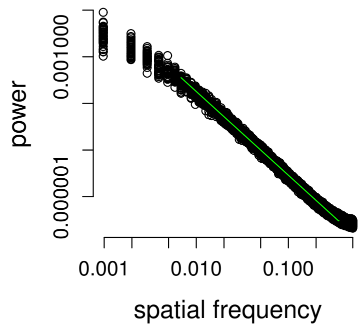


(d) $q = 0.92, p = 0.2852$, $PLR = 0.67$

Figure 5: The shift in the functional form of the power-spectrum is captured by the increased range of power-law as systems approach critical points. Data are shown with black dots and the fitted power-law functions, as green lines

To compare the range of power law (PLR), we use the method proposed by [1], who define it as:

$$PLR = 1 - \frac{log[x_{min}] - log[x_{smallest}]}{log[x_{max}] - log[x_{smallest}]}$$

PLR varies from 0 (when none of the data fall within the fitted power-law) to 1 (when all the data fall within the power-law region). We see in our power-spectra that as the system approaches a critical point, smaller and smaller spatial freqencies fall within the power-law range. This is congruent with theoretical predictions of diverging correlation length at critical points.

---
